## Supplementary Information for "Promiscuous NAD-dependent dehydrogenases enable efficient bacterial growth on the PET monomer ethylene glycol"

*[1] Institute of Biology Leiden, Leiden University, Leiden, The Netherlands; [2] Max Planck Institute for Terrestrial Microbiology, Marburg, Germany; [3] Netherlands Center for Electron Nanoscopy, Leiden University, Leiden, the Netherlands; [4] Department of Cell and Chemical Biology, Leiden University Medical Center, Leiden, The Netherlands; [5] Facility for Mass Spectrometry and Proteomics, Max Planck Institute for Terrestrial Microbiology, Marburg, Germany*

**Table of Contents Supplementary Information:**

| | |
|---|---|
| Supplementary Figures 1 – 8 | p. 2 – 9 |
| Supplementary Tables 1 – 5 | p. 10 – 13 |
| References | p. 14 |

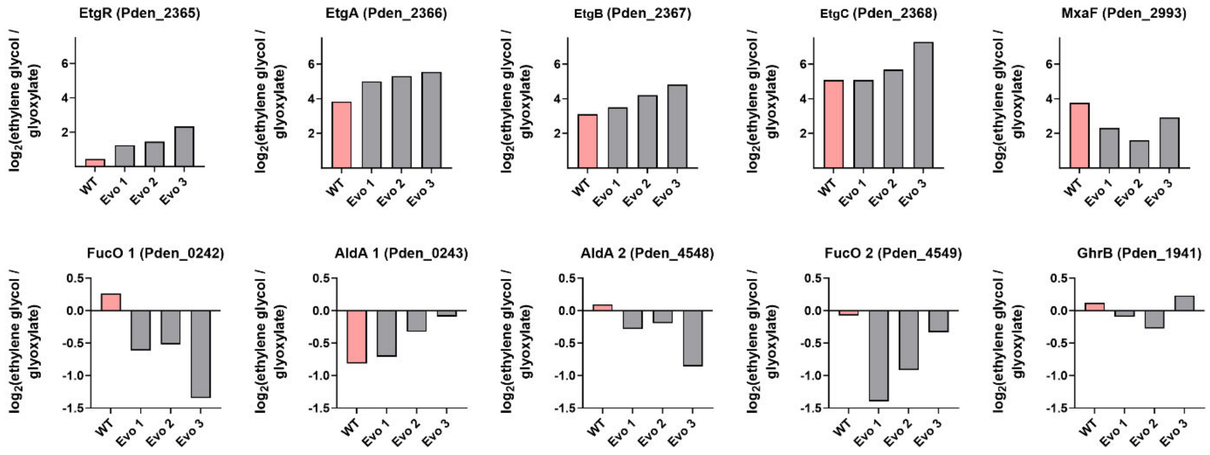


**Supplementary Figure 1: Proteome analysis demonstrates upregulation of the *etg* gene cluster during growth on ethylene glycol.** The $log_2$ fold change of EtgR/A/B/C in *P. denitrificans* WT and three evolved isolates growing on ethylene glycol compared to *P. denitrificans* WT growing on glyoxylate is shown. Similarly, the upregulation of MxaF is shown. In contrast, other alcohol and aldehyde dehydrogenases (FucO1/2, AldA1/2) as well as glyoxylate reductase (GhrB) either remained nearly unchanged or were downregulated.

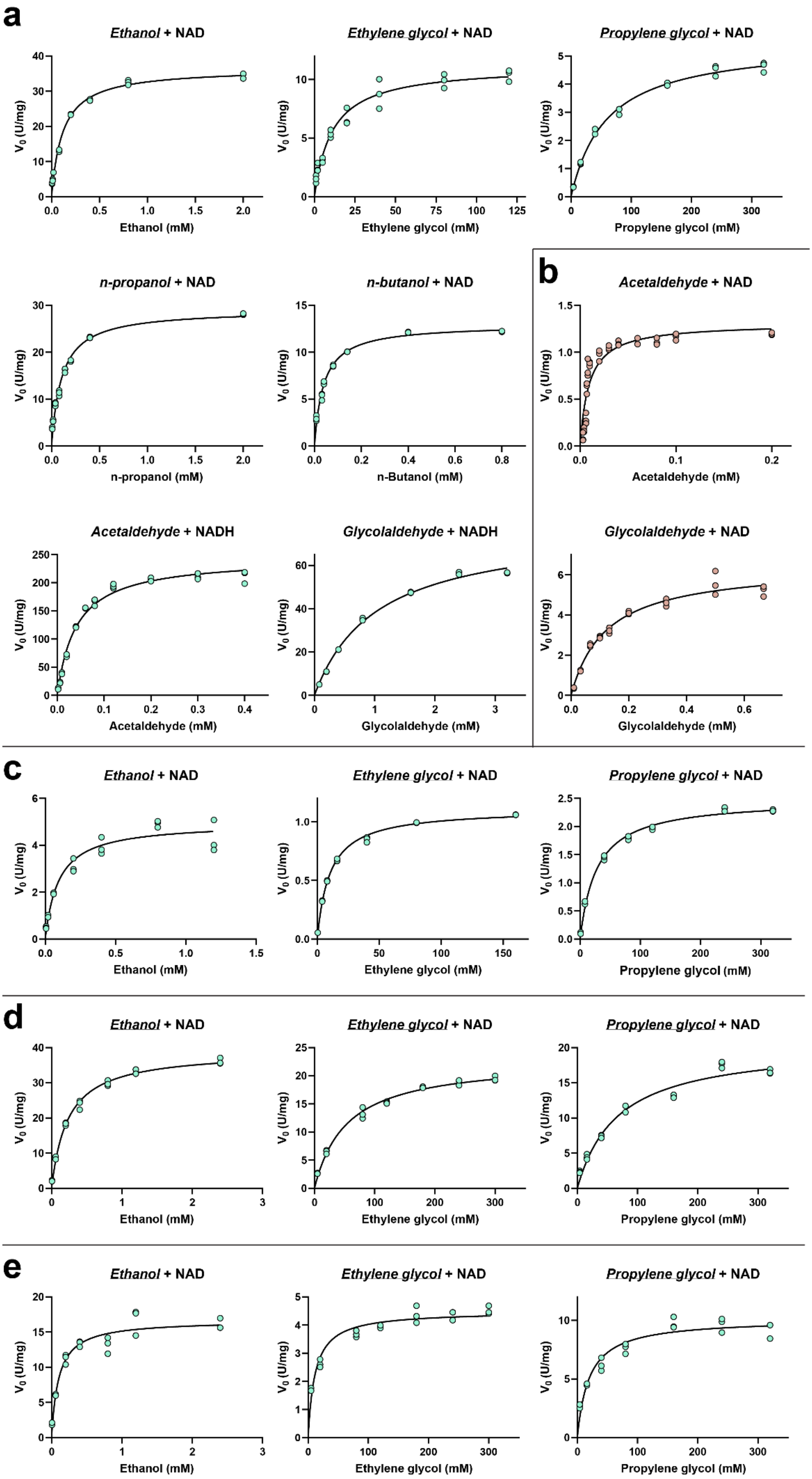

a
Ethanol + NAD
Ethylene glycol + NAD
Propylene glycol + NAD
Ethanol (mM)
Ethylene glycol (mM)
Propylene glycol (mM)
n-propanol + NAD
n-butanol + NAD
n-propanol (mM)
n-Butanol (mM)
b
Acetaldehyde + NAD
Acetaldehyde (mM)
Acetaldehyde + NADH
Glycolaldehyde + NADH
Glycolaldehyde + NAD
Acetaldehyde (mM)
Glycolaldehyde (mM)
Glycolaldehyde (mM)
c
Ethanol + NAD
Ethylene glycol + NAD
Propylene glycol + NAD
Ethanol (mM)
Ethylene glycol (mM)
Propylene glycol (mM)
d
Ethanol + NAD
Ethylene glycol + NAD
Propylene glycol + NAD
Ethanol (mM)
Ethylene glycol (mM)
Propylene glycol (mM)
e
Ethanol + NAD
Ethylene glycol + NAD
Propylene glycol + NAD
Ethanol (mM)
Ethylene glycol (mM)
Propylene glycol (mM)
V0 (U/mg)

**Supplementary Figure 2: Michaelis–Menten kinetics of all enzyme reactions characterized in this study. a**, Michaelis–Menten kinetics for EtgB. **b**, Michaelis–Menten kinetics for EtgA. **c**, Michaelis–Menten kinetics for EtgB T44S. **d**, Michaelis–Menten kinetics for EtgB H47N. **e**, Michaelis–Menten kinetics for EtgB T44S H47N. **a–e**, Data are shown from *n* = 3 independent experiments at different substrate concentrations. The data are summarized in **Table 1**.

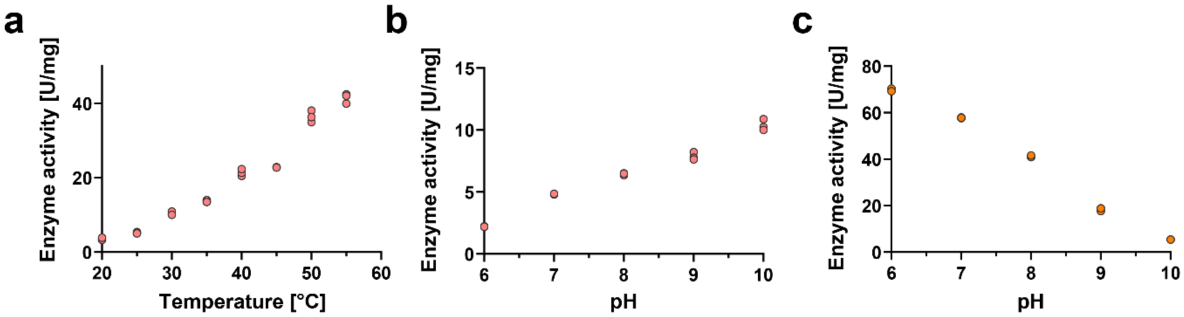


**Supplementary Figure 3: EtgB enzyme activity is dependent on temperature and pH. a**, Oxidation of ethylene glycol with $NAD^+$ as cofactor was measured at temperatures between 20 °C and 55 °C. **b**, Oxidation of ethylene glycol with $NAD^+$ as cofactor was measured at pH values from 6 to 10. **c**, Reduction of glycolaldehyde with NADH as cofactor was measured at pH values from 6 to 10. For **a** to **c**, the results of *n* = 3 independent experiments are shown.

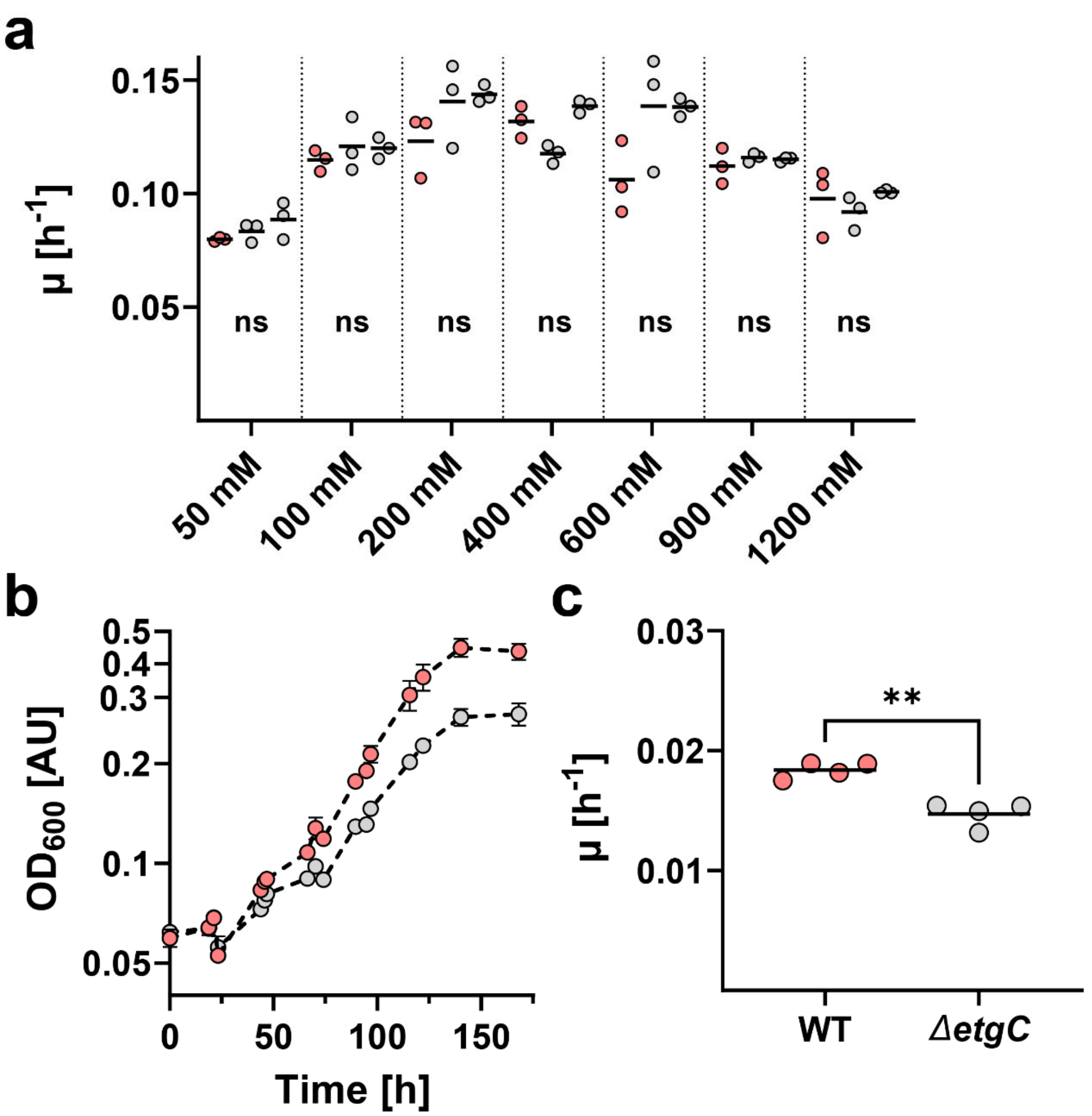


**Supplementary Figure 4: Growth of *P. denitrificans* Δ*etgC* under aerobic and anaerobic conditions.** **a**, Growth rates ($\mu$) of wild-type *P. denitrificans* DSM 413 (red) and *etgC* deletion strains (grey) grown aerobically in the presence of various concentrations of ethylene glycol. When compared to the wild-type, the growth rates of the Δ*etgC* strains were not significantly decreased. The results of $n = 3$ independent experiments are shown, and the black line represents the mean. **b**, Growth curves of the WT and Δ*etgC* strains in the presence of 30 mM ethylene glycol and 120 mM $KNO_3$ under anaerobic conditions. **c**, Growth rates calculated from this experiment. When compared to the wild-type, the growth rates of the Δ*etgC* strain were significantly decreased (p = 0.002). The results of $n = 4$ independent experiments are shown, and the black line represents the mean.

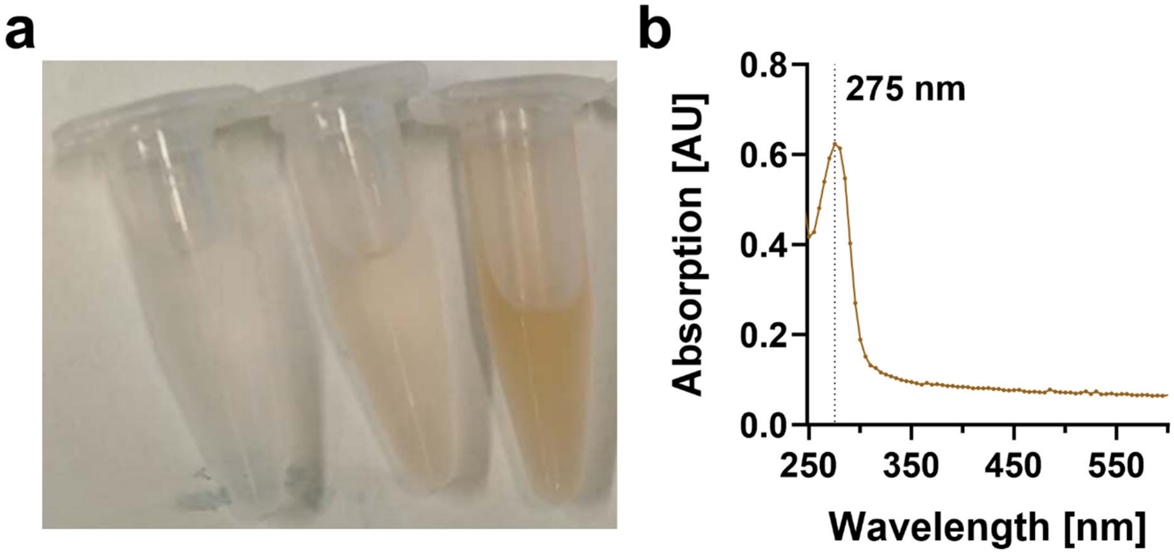


**Supplementary Figure 5: EtgC may contain an iron-sulfur cluster.** **a**, Increasing concentrations of aerobically purified EtgC (from left to right) exhibit a brownish color. **b**, Absorption spectrum of aerobically purified EtgC.

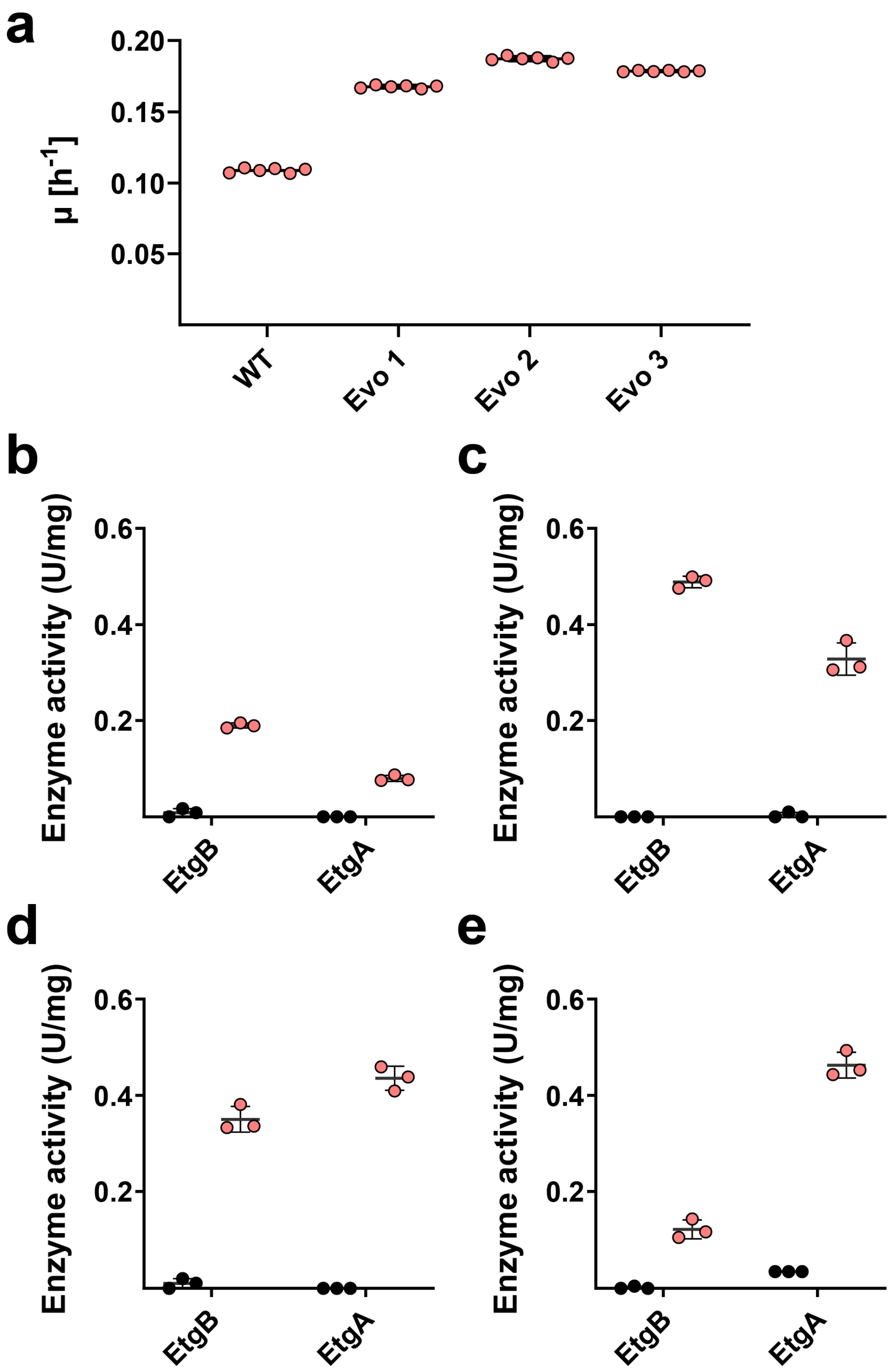


**Supplementary Figure 6: Characterization of *P. denitrificans* strains evolved for improved growth on ethylene glycol. a,** Growth rates of *P. denitrificans* WT and Evo1/2/3 on 200 mM ethylene glycol. The results of six independent experiments are shown. **b** to **e,** Cell-free extract enzyme assays for EtgA and EtgB. Specific activities of EtgB (with 200 mM ethylene glycol) and EtgA (with 0.6 mM glycolaldehyde) in cell-free extracts of *P. denitrificans* WT (**b**) and Evo1/2/3 (**c**, **d**, **e**, respectively) grown on 30 mM succinate (black) or 60 mM ethylene glycol (light red), as measured spectrophotometrically. Data are the mean ± s.d. of $n$ = 3 technical replicates.

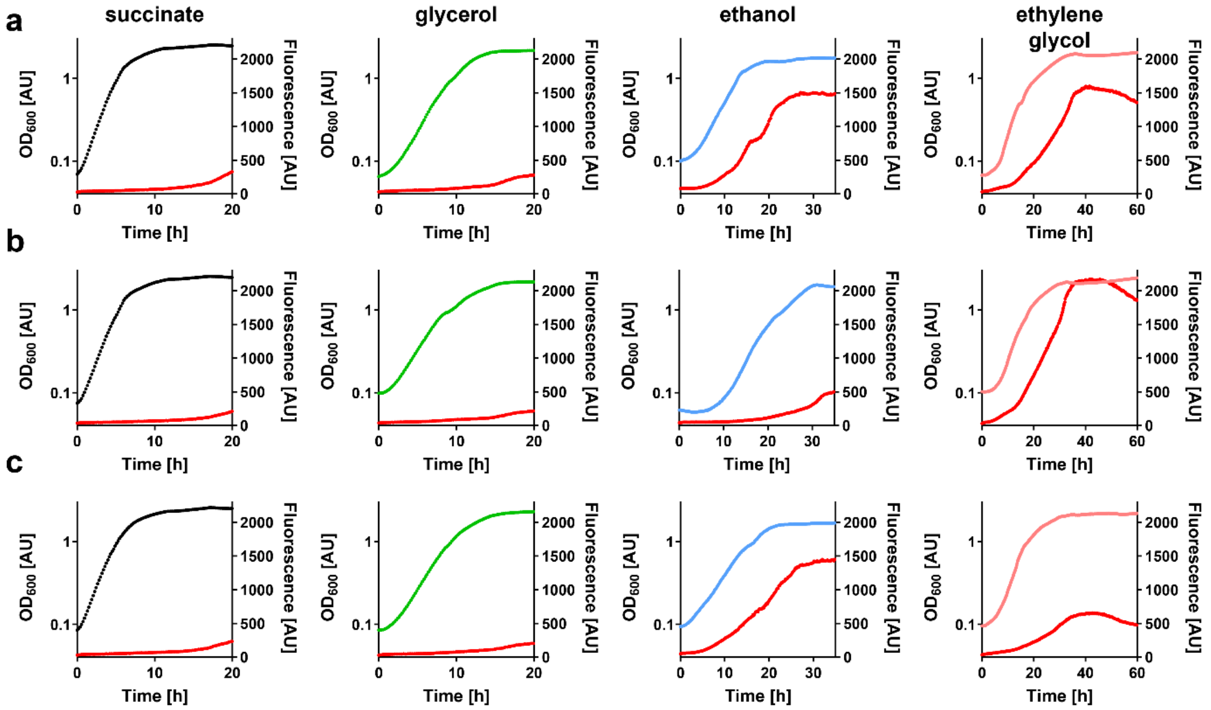


**Supplementary Figure 7: Characterization of evolved *P. denitrificans* promoter reporter strains on different carbon sources.** Growth and fluorescence (red) of *P. denitrificans* Evo 1 (**a**), Evo 2 (**b**), and Evo 3 (**c**) with pTE714-$P_{etg}$ on different carbon sources. These experiments were repeated three times independently with similar results. Growth and fluorescence of negative control strains are shown in **Supplementary Figure 8**.

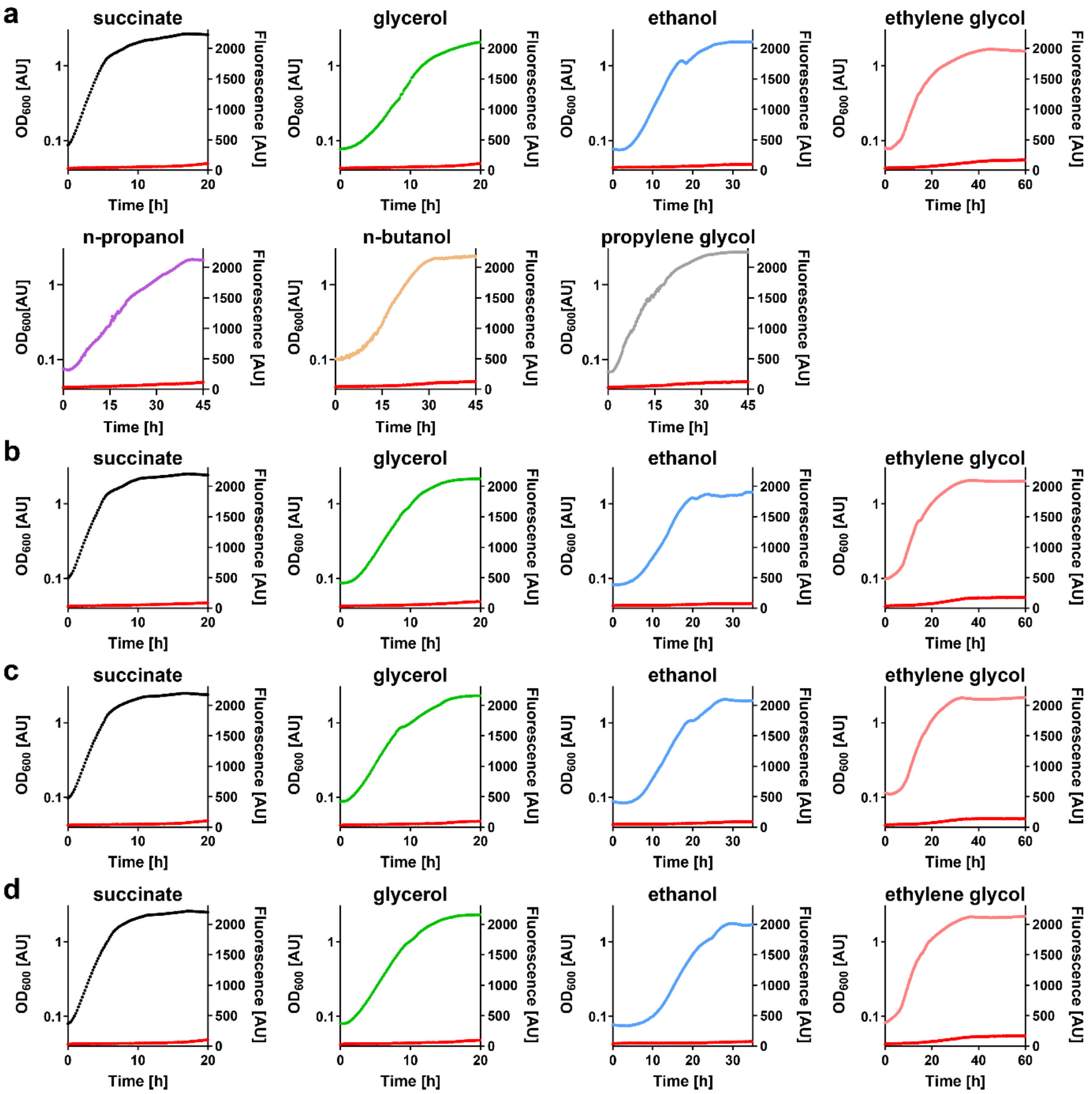


**Supplementary Figure 8: Characterization of *P. denitrificans* promoter reporter strains on different carbon sources (negative controls).** Growth and fluorescence (red) of *P. denitrificans* WT (**a**), Evo 1 (**b**), Evo 2 (**c**), and Evo 3 (**d**) with pTE714 (empty vector) on different carbon sources. These experiments were repeated three times independently with similar results.

**Supplementary Table 1: Kinetic parameters of NAD-dependent alcohol dehydrogenases with ethylene glycol as substrate.**

| **Enzyme** | **Organism** | $V_{max}$ **(U mg$^{-1}$)** | **App.** $K_m$ **(mM)** | $k_{cat}/K_m$ **($M^{-1}$ $s^{-1}$)** | **Assay conditions** | **Reference** |
|---|---|---|---|---|---|---|
| Alcohol dehydrogenase Gox0313 | *Gluconobacter oxydans* | 7.1 ± 0.3 | 964 ± 84 | 4.7 | 100 mM MOPS/KOH pH 7.8, 2 mM $NAD^+$, 37 °C | Scheffen et al. (1) |
| Alcohol dehydrogenase | *Pseudomonas aeruginosa* | 7.5 | > 200 | < 22.5 | 100 mM Tris-HCl pH 8.8, 0.5 mM $NAD^+$, 40 °C | Levin et al. (2) |
| Alcohol dehydrogenase | *Homo sapiens* | 1.1 | 290 | 2.6 | 100 mM glycine-NaOH pH 10, 2.4 mM $NAD^+$, 25 °C | Ditlow et al. (3) |
| Glycerol dehydrogenase | *Thermus thermophilus* | 30.3 | 56 | 328.4 | N/A | Raghava and Gupta (4) |
| Glycerol dehydrogenase | *Klebsiella pneumoniae* | 4.2 ± 0.3 | 92.7 ± 22.2 | 30.2 | 200 mM CHES pH 8.6, 10 mM $NAD^+$, 35 °C | Ko et al. (5) |
| Lactaldehyde reductase FucO | *Escherichia coli* | 5.9 ± 0.1 | 51 ± 2 | 80 | 100 mM glycine-NaOH pH 10, 0.2 mM $NAD^+$, 30 °C | Blikstad and Widersten (6) |

**Supplementary Table 2: Cryo-EM data collection, refinement and validation statistics.**

| | EtgA | EtgB |
|---|---|---|
| **Data collection** | | |
| Microscope | Titan Krios | Titan Krios |
| Voltage (kV) | 300 | 300 |
| Magnification | 105,000x | 105,000x |
| Detector - GIF | Gatan K3 - Bioquantum | Gatan K3 - Bioquantum |
| Data collection software | EPU | EPU |
| Electron exposure ($e^{-}/Å^{2}$) | 50 | 50 |
| Defocus range (µm) | 0.8 - 2.2 | 0.8 - 2.2 |
| Pixel size (Å) | 0.836 | 0.836 |
| **Data processing** | | |
| Software | Relion 5 | Relion 5 |
| Number of micrographs | 6697 | 5412 |
| Final number of particles | 185597 | 206685 |
| Symmetry imposed | C1 | C1 |
| Map resolution (Å) | 3.1 | 3.0 |
| FSC threshold | 0.143 | 0.143 |
| **Model Refinement** | | |
| Software | Phenix 1.18.2 | Phenix 1.18.2 |
| Map correlation coefficient | 0.78 | 0.79 |
| Model composition | | |
| Number of chains | 4 | 4 |
| Non-hydrogen atoms | 12756 | 9910 |
| Protein residues | 1654 | 1346 |
| Ligands | 2 | 8 |
| R.M.S. deviations | | |
| Bond lengths (Å) | 0.006 | 0.008 |
| Bond angles (°) | 0.720 | 0.835 |
| **Validation** | | |
| MolProbity score | 2.12 | 2.07 |
| Clashscore | 14.5 | 15.06 |
| Rotamer outliers (%) | 0 | 0 |
| Ramachandran plot | | |
| Favored (%) | 92.98 | 94.29 |
| Allowed (%) | 7.02 | 5.71 |
| Disallowed (%) | 0 | 0 |
| **Data availability** | | |
| EMDB entry | EMD-50550 | EMD-50545 |
| PDB entry | 9FM9 | 9FLZ |

### Supplementary Table 3: Strains used in this study

| strain | genotype or relevant features[a] | source or reference |
|---|---|---|
| *E. coli* DH5α | *supE44*, *ΔlacU169* (*Φ80lacZDM15*), *hsdR17*, *recA1*, *endA1*, *gyrA96*, *thi*-1, *relA1* | Thermo Fisher Scientific, Waltham, USA |
| *E. coli* ST18 | *pro*, *thi*, *hsdR1*,Tp$^R$, Sm$^R$; chromosome::RP4-2, Tc::Mu-Kan::Tn7/λpir, λpir, Δ*hemA* | Thoma and Schobert (7) |
| *E. coli* BL21 AI | *ompT*, *gal*, *dcm*, *lon*, $hsdS_B(r_B^- m_B^-)$, $[malB^+]_{K-12}(\lambda^S)$, *araB*::*T7RNAP-tetA* | Thermo Fisher Scientific, Waltham, USA |
| *P. denitrificans* DSM 413 | WT strain | Beijerinck and Minkman (8) |
| *P. denitrificans* DSM 413 Δ*etgR* 1 | Δ*etgR*; Km$^R$ (orientation 1) | this work |
| *P. denitrificans* DSM 413 Δ*etgR* 2 | Δ*etgR*; Km$^R$ (orientation 2) | this work |
| *P. denitrificans* DSM 413 Δ*etgC* 1 | Δ*etgC*; Km$^R$ (orientation 1) | this work |
| *P. denitrificans* DSM 413 Δ*etgC* 2 | Δ*etgC*; Km$^R$ (orientation 2) | this work |
| *P. denitrificans* DSM 413 Evo 1 | evolved on ethylene glycol; see Table 2 | this work |
| *P. denitrificans* DSM 413 Evo 2 | evolved on ethylene glycol; see Table 2 | this work |
| *P. denitrificans* DSM 413 Evo 3 | evolved on ethylene glycol; see Table 2 | this work |

[a] Km$^R$, kanamycin resistance; Tp$^R$, trimethoprim resistance; Sm$^R$, streptomycin resistance

### Supplementary Table 4 | Plasmids used in this study

| plasmid | relevant features[a] | source or reference |
|---|---|---|
| pET16b | *E. coli* expression vector, T7 promoter, Amp$^R$ | Merck Chemicals GmbH, Darmstadt, Germany |
| pET16b-EtgA | expression vector for N-terminally His-tagged EtgA, Amp$^R$ | this work |
| pET16b-EtgB | expression vector for N-terminally His-tagged EtgB, Amp$^R$ | this work |
| pET16b-EtgB T44S | expression vector for N-terminally His-tagged EtgB T44S, Amp$^R$ | this work |
| pET16b-EtgB H47N | expression vector for N-terminally His-tagged EtgB H47N, Amp$^R$ | this work |
| pET16b-EtgB T44S H47N | expression vector for N-terminally His-tagged EtgB T44S H47N, Amp$^R$ | this work |
| pET16b-EtgC | expression vector for N-terminally His-tagged EtgC, Amp$^R$ | this work |
| pREDSIX | mobilizable, high-copy-number cloning and mutagenesis vector; Amp$^R$ | Ledermann et al. (9) |
| pRGD-KmR | donor vector for resistance gene, *aphII* in polylinker; Amp$^R$, Km$^R$ | Ledermann et al. (9) |
| pREDSIX-*etgR-1* | knockout vector for the *etgR* gene in *P. denitrificans* DSM 413; Km$^R$ (orientation 1) | this work |
| pREDSIX-*etgR-2* | knockout vector for the *etgR* gene in *P. denitrificans* DSM 413; Km$^R$ (orientation 2) | this work |
| pREDSIX-*etgC-1* | knockout vector for the *etgC* gene in *P. denitrificans* DSM 413; Km$^R$ (orientation 1) | this work |
| pREDSIX-*etgC-2* | knockout vector for the *etgC* gene in *P. denitrificans* DSM 413; Km$^R$ (orientation 2) | this work |
| pTE714 | promoter probe vector for Alphaproteobacteria containing a RBS and mCherry; Tc$^R$ | Schada von Borzyskowski et al. (10) |
| pTE714_2365/2366_ig | promoter probe vector containing mCherry under control of the promoter located in the intergenic region between Pden_2365 and Pden_2366; Tc$^R$ | this work |

[a] Km$^R$, kanamycin resistance; Amp$^R$, ampicillin resistance; Tc$^R$, tetracycline resistance

### Supplementary Table 5 | Primers used in this study

| target | name | sequence[a] | cut site |
|---|---|---|---|
| Pden_*etgA* | etgA_16b_fw | 5'-CTAGAGT**CATATG**CCGAACGACCAGACG-3' | *Nde*I |
| Pden_ *etgA* | etgA_16b_rv | 5'-GACTACT**GGATCC**TCAGAAGAAGCCCAGCTTC-3' | *Bam*HI |
| Pden_ *etgB* | etgB_16b_fw | 5'-CTCGTAC**ATTAAT**ATGGCCAAAACCATGAAAGCCG-3' | *Ase*I |
| Pden_ *etgB* | etgB_16b_rv | 5'-CATTACT**CTCGAG**TCAGCCCGCCATGTCCAGCAC-3' | *Xho*I |
| Pden_ *etgC* | etgC_16b_fw | 5'-GACTAG**CATATG**GAACCCGTCGCCACCC-3' | *Nde*I |
| Pden_ *etgC* | etgC_16b_rv | 5'-GTTTAC**GGATCC**TCAGAGCCGGCAGATCTCGGAC-3' | *Bam*HI |
| Pden_ *etgB* | T44S_fw | 5'-TGCCATAGCGACCTGCACGCGGCCGAGGGC-3' | --- |
| Pden_ *etgB* | T44S_rv | 5'-GGTCGCTATGGCAGACGCCCGAGGCCTG-3' | --- |
| Pden_ *etgB* | H47N_fw | 5'-ACCTGAACGCGGCCGAGGGCGACTG-3' | --- |
| Pden_ *etgB* | H47N_rv | 5'-CGGCCGCGTTCAGGTCGGTATGGCAGA-3' | --- |
| Pden_ *etgB* | T44S_H47N_fw | 5'-GCCATTCGGACCTGAACGCGGCC-3' | --- |
| Pden_ *etgB* | T44S_H47N_rv | 5'-TCAGGTCCGAATGGCAGACGCCCG-3' | --- |
| Pden_*etgR*_up | etgR_up_fw | 5'-GGTCTGACAGGTTTAAACTCTAGACGGATCGCCGAGCGCATCGCGC-3' | --- |
| Pden_*etgR*_up | etgR_up_rv | 5'-GCCCCGATGTGTT**CATATG**CGAAGCACCCCGGCCTTTC-3' | *Nde*I |
| Pden_*etgR*_down | etgR_down_fw | 5'-GCTTCG**CATATG**AACACATCGGGGCAGCGGCGCG-3' | *Nde*I |
| Pden_*etgR*_down | etgR_down_rv | 5'-CTTAAGGCTAGCATGCATCCTAGGCAGCACGCCCGGCGGCAGCAGG-3' | --- |
| Pden_*etgC*_up | etgC_up_fw | 5'-GGTCTGACAGGTTTAAACTCTAGACCGGCCATTGCCGGCATTGCCT-3' | --- |
| Pden_*etgC*_up | etgC_up_rv | 5'-CCCGTGCCGCGG**GGTACC**GCTCAGCCCGCCATG-3' | *Kpn*I |
| Pden_*etgC*_down | etgC_down_fw | 5'-GCTGAGC**GGTACC**CCGCGGCACGGGCGCACGGCC-3' | *Kpn*I |
| Pden_*etgC*_down | etgC_down_rv | 5'CTTAAGGCTAGCATGCATCCTAGGCTGAAGACCCCCTCGCGGCTGATCGGC ATGATCCAG-3' | --- |
| Pden_2365/2366_ig | 2365/2366_ig_fw | 5'-TTGACA**GAATTC**CGAGGGCAATGTCTCGGC-3' | *Eco*RI |
| Pden_2365/2366_ig | 2365/2366_ig_rv | 5'-ACTCAAT**TCTAGA**TCAGATGTTCCTCCTGTCATTCCGGCT-3' | *Xba*I |

[a] Nucleotides in bold and underlined are recognition sites for endonuclease restriction enzymes.
